# The Spx stress regulator confers high-level β-lactam resistance and decreases susceptibility to last-line antibiotics in methicillin resistant *Staphylococcus aureus*

**DOI:** 10.1101/2024.03.01.582999

**Authors:** Tobias Krogh Nielsen, Ida Birkjær Petersen, Lijuan Xu, Maria Disen Barbuti, Viktor Mebus, Anni Justh, Abdulelah Ahmed Alqarzaee, Nicolas Jacques, Cécile Oury, Vinai Thomas, Morten Kjos, Camilla Henriksen, Dorte Frees

## Abstract

Infections caused by methicillin resistant *Staphylococcus aureus* (MRSA) are a leading cause of mortality worldwide. MRSA have acquired resistance to next generation β-lactam antibiotics through the horizontal acquisition of the *mecA* resistance gene. Development of high resistance is, however, often associated with additional mutations in a set of chromosomal core genes, known as potentiators which through poorly described mechanisms enhance resistance. The *yjbH* gene was recently identified as a hot spot for adaptive mutations during severe infections. Here, we show that inactivation of *yjbH* increased β-lactam MICs up to 16-folds and transformed MRSA cells with low level of resistance to being homogenously highly resistant to β-lactams. The *yjbH* gene encodes an adaptor protein that targets the transcriptional stress regulator Spx for degradation by the ClpXP protease. Using CRISPRi to knock down *spx* transcription, we unambiguously linked hyper-resistance to accumulation of Spx. Spx was previously proposed to be essential, however, our data indicate that Spx is dispensable for growth at 37°C but becomes essential in the presence of antibiotics with various targets. On the other hand, high Spx levels bypassed the role of PBP4 in β-lactam resistance and broadly decreased MRSA susceptibility to compounds targeting the cell wall or the cell membrane including vancomycin, daptomycin, and nisin. Strikingly, Spx potentiated resistance independently of its redox sensing switch. Collectively, our study identifies a general stress pathway that, in addition to promoting the development of high-level, broad-spectrum β-lactam resistance, also decreases MRSA susceptibility to critical antibiotics of last resort.

## Introduction

*Staphylococcus aureus* is a natural part of the human microbiome and colonizes approximately 30% of the healthy human population (1,2). Also, *S. aureus* is notorious for its ability to transform into an aggressive pathogen that in hospitalized, immunocompromised patients is a leading cause of deadly infections such as bacteremia, sepsis, osteomyelitis, and infective endocarditis (2,3). Historically, β-lactam antibiotics have been the agents of choice for the treatment of staphylococcal infections but treatment is hampered by the world-wide dissemination of methicillin-resistant *S. aureus* (MRSA) that through the horizontal acquisition of the resistance gene, *mecA*, have developed resistance to virtually all β-lactam antibiotics (4,5). Despite that antibiotic resistance is often portrayed as a black-and-white phenomenon, the *mecA* resistance gene often only confers low-level β-lactam resistance with some *mecA*-positive *S. aureus* being phenotypically sensitive to β-lactams (6–9). The development of high-level resistance requires additional potentiating mutations in a restricted set of chromosomal core genes not linked to the cell wall target of the antibiotic (10–12). So far, the molecular pathways enabling potentiators to profoundly increase resistance remain poorly described as elegantly summarized in a recent review (11).

All β-lactam antibiotics share a common mechanism of action involving the irreversible binding and inactivation of a class of enzymes, the so-called penicillin-binding proteins (PBPs) that catalyze the cross-linking of cell wall peptidoglycan (12). *S. aureus* encodes four native PBPs (PBP1-4) of which only PBP1 and PBP2 are essential (13). The *mecA* gene encodes an alternative transpeptidase, PBP2a that has low affinity for β-lactams allowing it to perform the critical cross-linking when native PBPs are inhibited by the irreversible binding of β-lactams (14). Curiously, *mecA* mediated resistance to β-lactams is typically expressed in a heterogeneous fashion with the majority of the bacterial population being sensitive to concentrations of antibiotic far below the MIC, while only a small subpopulation, typically 0.01–0.1%, displays higher levels of resistance (15,16). The development of homogenous, high-level resistance is associated with potentiating mutations in a restricted set of chromosomal core genes encoding *rpoB* and *rpoC* genes encoding the β and β′ subunits of the RNA polymerase, the ClpXP protease, and genes controlling synthesis of the nucleotide-signaling molecules c-di-AMP and (p)ppGpp (10–12; 17–22).

In Firmicutes, Spx is a highly conserved transcriptional regulator that is best known for its role in controlling a disulfide stress response (23–25). Non-stressed cells have low Spx levels because the YjbH adaptor continuously targets Spx for degradation by the ClpXP protease composed of separately encoded proteolytic subunits (ClpP) and unfolding subunits (ClpX) (26,27). In *S. aureus*, a comprehensive genetic study identified the *yjbH* gene as a hot spot for adaptive mutations during severe infections (28). Further mutations in *yjbH* have been associated with non-*mec*-mediated oxacillin resistance that was tentatively explained by overproduction of the non-essential transpeptidase PBP4, a critical determinant of MRSA resistance (29–34). Here, we identify the *yjbH* gene as a potentiator of β-lactam resistance also in MRSA. By using CRISPR interference (CRISPRi) knock-down of *spx* and *pbp4* transcription, we unambiguously link the β-lactam hyper resistance phenotype of MRSA *yjbH* mutants to the Spx transcriptional regulator and show that PBP4 becomes dispensable for resistance in cells with high Spx. Spx broadly promoted growth of MRSA in the presence of compounds targeting the cell wall or the cell membrane independently of its redox sensing switch. Collectively, the data presented support that Spx has a pivotal role in protecting MRSA against β-lactams and other critical antibiotics targeting the cell envelope.

## Results

### Inactivation of *yjbH* enhances β-lactam resistance in MRSA USA300

Inactivating mutations in the *yjbH* gene have been shown to play a role in non-*mecA* mediated resistance (29–32), and we here investigated if inactivation of the *yjbH* gene also potentiates resistance in MRSA. As model strain we used the highly virulent, community-acquired USA300 clone that is characterized by expressing a relatively low level of resistance (12,33). In broth microdilution assays, a profound increase in β-lactam MICs was observed for the *yjbH* transposon mutant (35) as compared to the parental USA300-JE2 MRSA model strain (Table 1). In particular, the inactivation of *yjbH* resulted in a 16-fold increase in resistance to meropenem and a > 8-fold increase in oxacillin- and imipenem MICs (Table 1). Resistance was additionally investigated at the population level by performing population analysis profiles (PAPs). In PAP, the JE2 wild-type strain displayed typical heterogeneous resistance to β-lactams, however, inactivation of *yjbH* transformed the JE2 strain into being homogeneously, highly resistant to oxacillin (Fig. 1A). JE2 is cured of the conserved plasmid encoding the resistance gene *blaZ* along with the *blaR1*/*blaI* regulatory system (35–37). Therefore, we also confirmed the impact of the *yjbH* gene on resistance in the plasmid containing MRSA USA300 reference strain BAA(Fig. 1A). To investigate if the cellular amount of PBP2a was affected by disruption of *yjbH*, Western blotting was performed on total cellular extracts derived from BAA and JE2 wild-types and *yjbH* mutants grown with or without oxacillin (Fig. 1B). In the JE2 strain background, a small up-regulation of PBP2A was observed in the *yjbH* mutant, however, a similar upregulation was not observed in the BAA background (Fig. 1B). We conclude that inactivation of the *yjbH* gene greatly potentiates β-lactam resistance in the clinically important USA300 MRSA clone via a mechanism that does not seem to depend on an up-regulation of PBP2a expression.

**Fig. 1.**
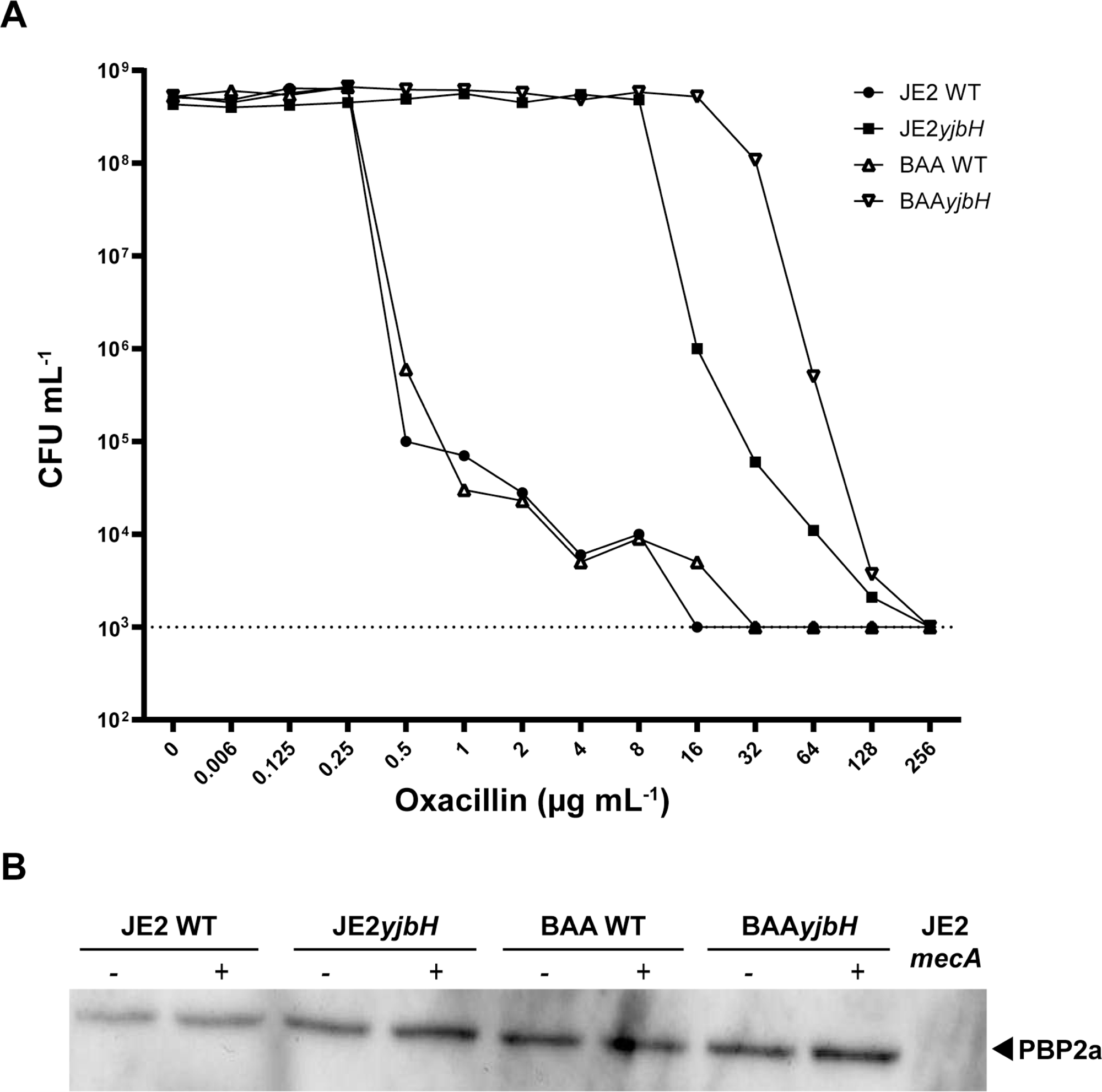
Inactivation of *yjbH* renders MRSA homogenously, highly resistant to oxacillin without changing PBP2a expression. (A) CFU ml^-1^ was determined after plating on increasing concentrations of oxacillin as indicated. Representative data from three individual experiments are shown. The dashed line indicates detection limit. (B) Cellular concentration of PBP2a in the BAA and JE2 strain background and their corresponding *yjbH* mutants was determined by Western blotting in the absence (-) or 30 minutes following exposure to 2 µg mL^-1^ oxacillin. The JE2 *mecA* mutant was included as a negative control.

**Table 1.** Antibiotic susceptibility of USA300-JE2 and the corresponding *yjbH* derivative to seven different antibiotics.

| MIC ($\mu\text{g ml}^{-1}$) | JE2 WT | JE2*yjbH* |
| --- | --- | --- |
| Oxacillin | 32 | >256 |
| Imipenem | 0.12 | 1 |
| Cefotaxime | 16 | >64 |
| Cefoxitin | 32 | >64 |
| Cefepime | 16 | >32 |
| Ertapenem | 2 | 2 |
| Meropenem | 0.5 | 8 |

### Elevated PBP4 is not causing the increase in resistance

In methicillin sensitive *S. aureus*, inactivation of *yjbH* gene was reported to cause a modest increase in resistance to β-lactams that was tentatively attributed to slightly increased amounts of PBP4 and enhanced cross-linking of peptidoglycan strands (29). Consistent with this finding, we observed that PBP4 levels were slightly enhanced upon inactivation of *yjbH* in the JE2 strain background (Fig. 2A). To investigate if the elevated PBP4 levels were causing increased resistance to β-lactams, we performed CRISPRi knockdown of the *pbp4* gene using the two plasmids system described by Stamsås et al. (38). In this system, a gene-specific single guide RNA (sgRNA) is constitutively expressed from one plasmid while the catalytically inactivated dCas9 is expressed from an IPTG inducible promoter, allowing for depletion of proteins by a gene-specific block in transcription when cells are grown with IPTG (Supplemental. Fig. 1) (38). Using Western blotting we verified a profound reduction of PBP4 in cells expressing dCAS9 together with an sgRNA targeting *pbp4*, while control cells expressing dCAS9 and a non-targeting sgRNA did not alter PBP4 levels (Fig. 2A). Knockdown of PBP4 expression did not reduce growth in either the wild-type nor in the *yjbH* mutant (Fig. 2B). We then determined oxacillin MICs after *pbp4* knockdown in JE2 wild-type and the JE2 *yjbH* mutant (see Supplemental Figure 1 for set-up of MIC assays). In agreement with published data showing that PBP4 is essential for β-lactam resistance in the USA300 (33,34), knockdown of *pbp4* transcription reduced the oxacillin MIC of JE2 from 32 µg mL^-1^ to 4 µg mL^-1^ while there was no effect of the nontargeting sgRNA (Table 2A). Strikingly, there was no effect of depleting PBP4 in the hyper-resistant *yjbH* mutant, showing that PBP4 is not contributing to the increased β-lactam resistance of *S. aureus* cells devoid of YjbH activity and that inactivation of *yjbH* even bypasses the need for PBP4 to achieve the wild-type resistance level (Table 2A).

**Fig. 2.**
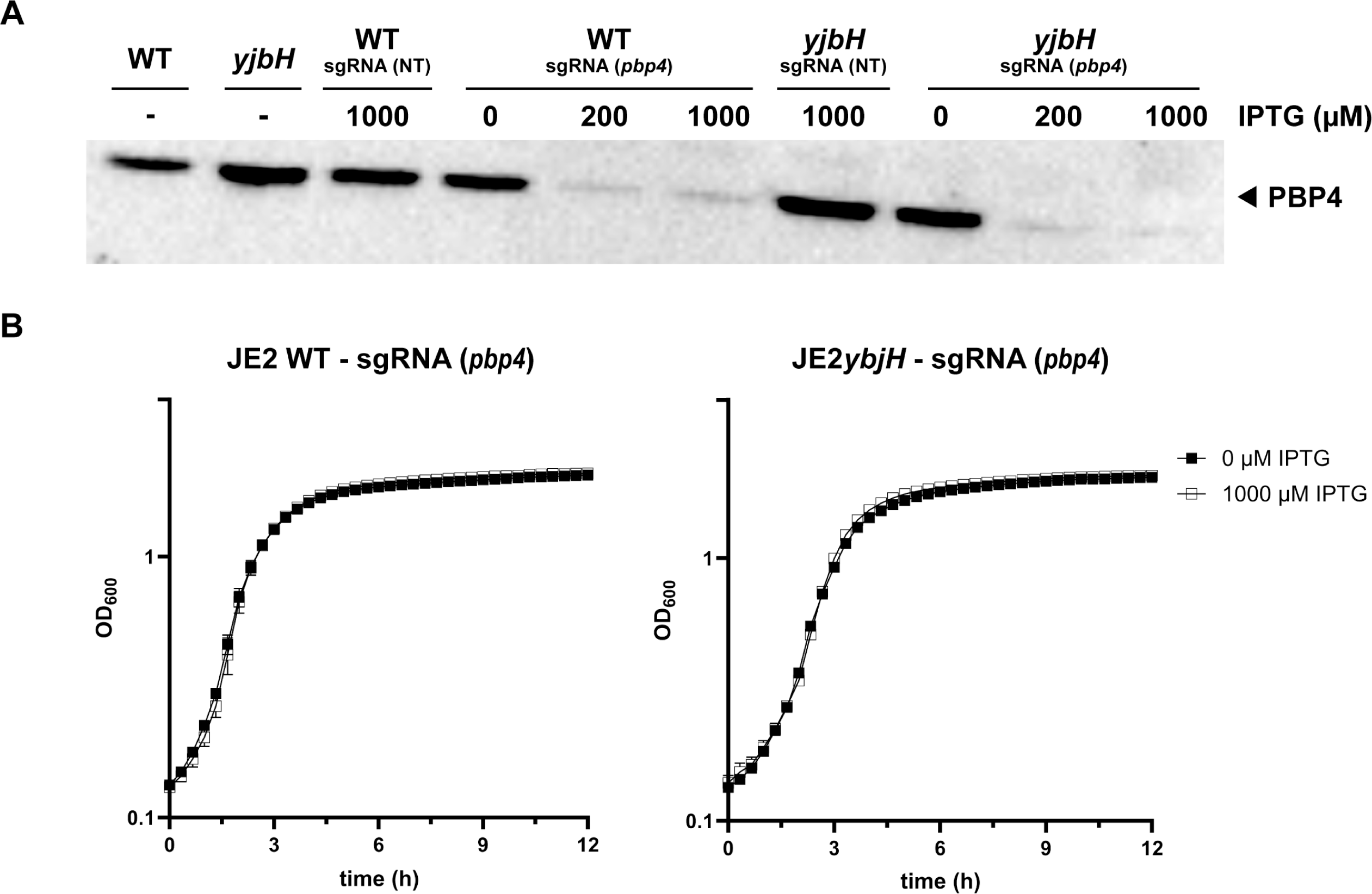
PBP4 is not causing the increase in resistance. (A) Western blot showing cellular concentration of PBP4 in the JE2 WT, JE2 *yjbH* mutant, and in strains harboring the two plasmids allowing CRISPRi knock down of *pbp4* transcription. Cells were harvested under non-inducing condition (no IPTG) and 30 min after inducing dCas9 expression with IPTG as indicated. Strains expressing a non-targeting (NT) sgRNA were included as controls. (B) Growth at 37°C for JE2 WT and JE2 *yjbH* mutant harboring the two plasmids allowing CRISPRi knockdown of *pbp4* transcription. Strains were grown in the absence or presence of 1000 µM IPTG at 37°C for 12 hours.

**Table 2.**
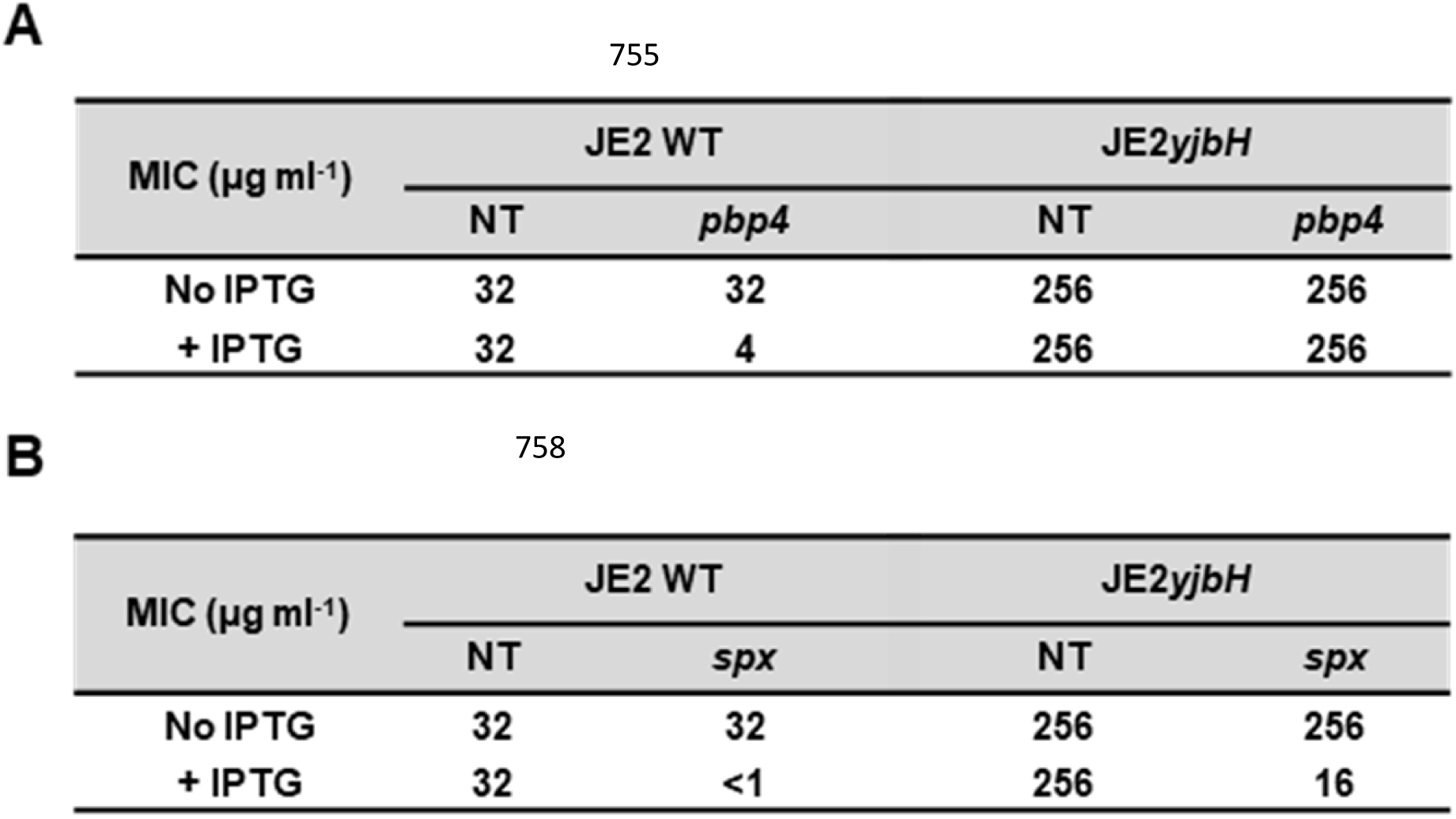
Oxacillin susceptibility of USA300-JE2 strain and the corresponding *yjbH* derivative with uninduced or induced CRISPRi transcription knockdown of *pbp4* (A) or *spx* (B).

### Resistance correlates positively to Spx levels

So far, the only known role of *S. aureus* YjbH is its role in targeting the transcriptional stress regulator Spx for degradation by the ClpXP protease (27). Accordingly, a Western blot confirmed that Spx accumulates in MRSA cells with inactivated YjbH (Fig. 3A). To examine if the accumulation of Spx contributes to increased resistance, we designed an sgRNA targeting *spx*, and *spx* knockdown by CRISPRi proved to be successful in reducing Spx levels below the detection level (Fig. 3A; Supplemental Fig. 1). The *spx* gene was in previous studies suggested to be essential for growth of *S. aureus* (39). Therefore, we were surprised to see that cells with *spx*-knockdown were able to form colonies (Fig. 3B). However, the JE2*yjbH* colonies, that are normally non-pigmented, regained pigmentation upon induction of the CRISPRi system (Fig. 3B). The increased pigmentation is supportive of successful *spx* knock-down as Spx is known to down-regulate synthesis of staphyloxanthin, the carotenoid causing the golden pigmentation of *S. aureus* colonies (40). Depletion of Spx did not compromise growth in liquid cultures, however, in spot-dilution assays, cells expressing the *spx*-specific sgRNA formed smaller colonies than CRISPRi control cells carrying a non-targeting sgRNA showing that Spx is indeed important for normal growth of *S. aureus* (Fig. 3C and 3D). Most interestingly, Spx depletion severely compromised the plating efficiencies of wild-type and *yjbH* cells in the presence of 0.05 µg mL^-1^ oxacillin (Fig. 3D). Consistent, with the spot-dilution assays, oxacillin MICs dropped from 32 µg ml^-1^ to below 1 µg ml^-1^ when Spx was depleted in the JE2 parental strain, and from 256 to 16 µg ml^-1^ in for the JE2*yjbH* mutant strain (Table 2B). Hence, the oxacillin MICs of JE2*yjbH* were always higher than in wild-type cells indicating that the block in YjbH dependent proteolysis of Spx results in higher Spx levels also under conditions where *spx* transcription is blocked by the CRISPRi system. To investigate if resistance correlated to Spx levels, we utilized the titratable CRISPRi-system to set up a checker-board assay where oxacillin MICs were determined in the presence of two-fold serial dilutions of IPTG (from 0 to 1 mg ml^-1^, higher IPTG concentrations resulting in more efficient depletion of Spx). Importantly, oxacillin MICs increased with decreasing IPTG concentrations (i.e. increasing Spx levels) in both JE2 WT and the JE2*yjbH* mutant, while oxacillin MICs remained constant in CRISPRi control cells strains expressing the non-targeting RNA (Fig. 4). We conclude that resistance levels are correlating positively with Spx abundance.

**Fig. 3.**
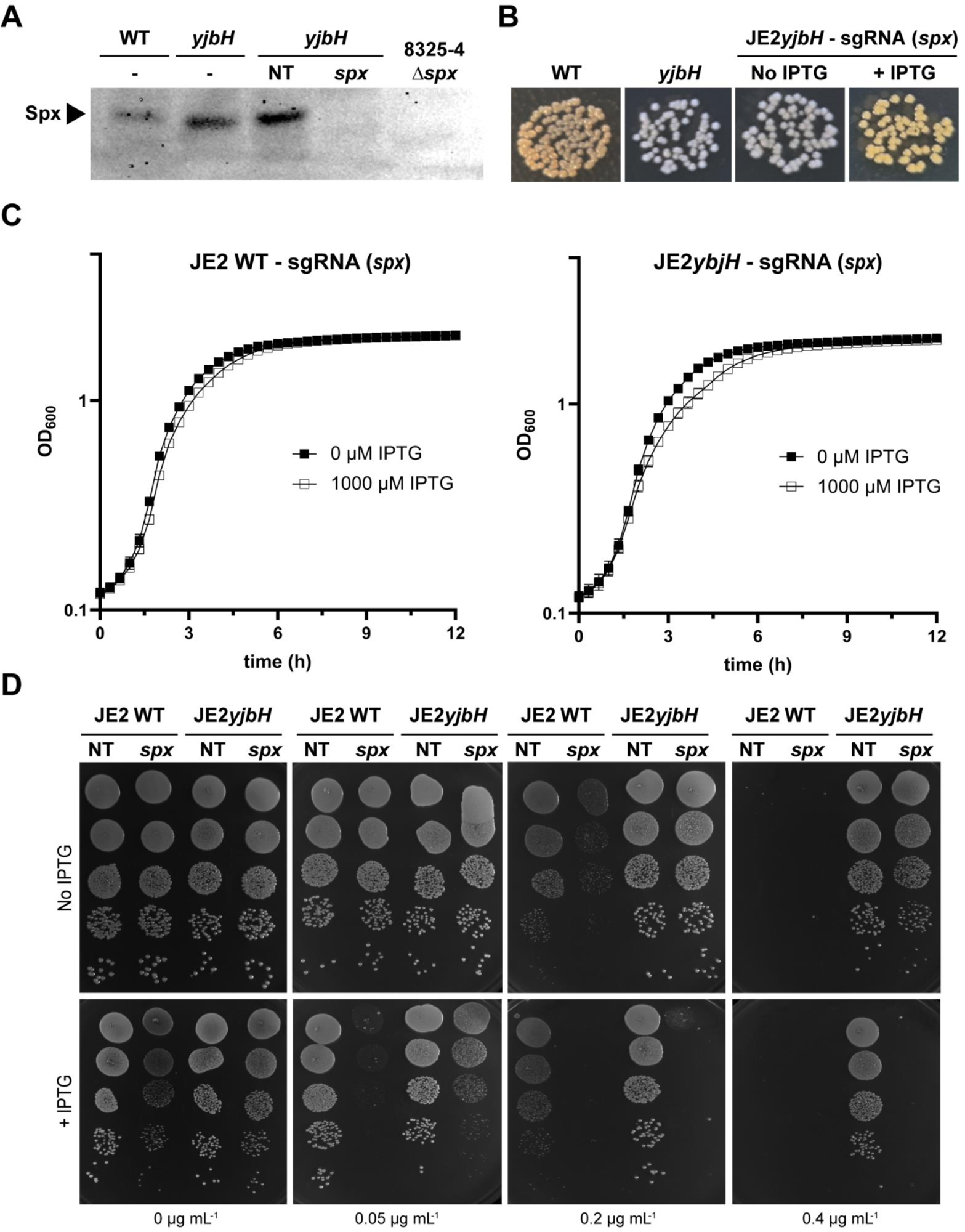
Spx depleted cells form colonies of increased pigmentation in the absence of antibiotics but cannot form colonies in the presence of oxacillin. (A) Western blot showing cellular Spx in over-night cultures of JE2 WT, the JE2 *yjbH* mutant, and in JE2 *yjbH* harboring the two plasmids allowing CRISPRi knock down of *spx* transcription. The CRISPRi-strain was grown in the absence or presence of IPTG as indicated. JE2 *yjbH* expressing a non-targeting (NT) sgRNA was included as negative control. A representative blot from two individual experiments is shown. (B) Pigmentation of colonies of the indicated strains spotted in 10 μL aliquots of on TSA ± 1000 µM IPTG as indicated and incubated at 37°C for 24 hours. (C) Growth curves of JE2 WT and the JE2 *yjbH* mutant (both strains harbor the two plasmids allowing CRISPRi knockdown of *spx* transcription) following growth at 37°C in TSB in the absence or presence of IPTG as indicated. Data represent three biological replicates. (D) Serial dilutions of the JE2 wild-type and JE2 *yjbH* mutant (both strains harbor the two plasmids allowing CRISPRi knockdown of *spx* transcription) were spotted in 10 μL aliquots on TSA ± 1000 µM IPTG and with increasing concentrations of oxacillin as indicated. JE2 wild-type and JE2 *yjbH* mutant expressing a non-targeting (NT) sgRNA were included as controls

**Fig. 4.**
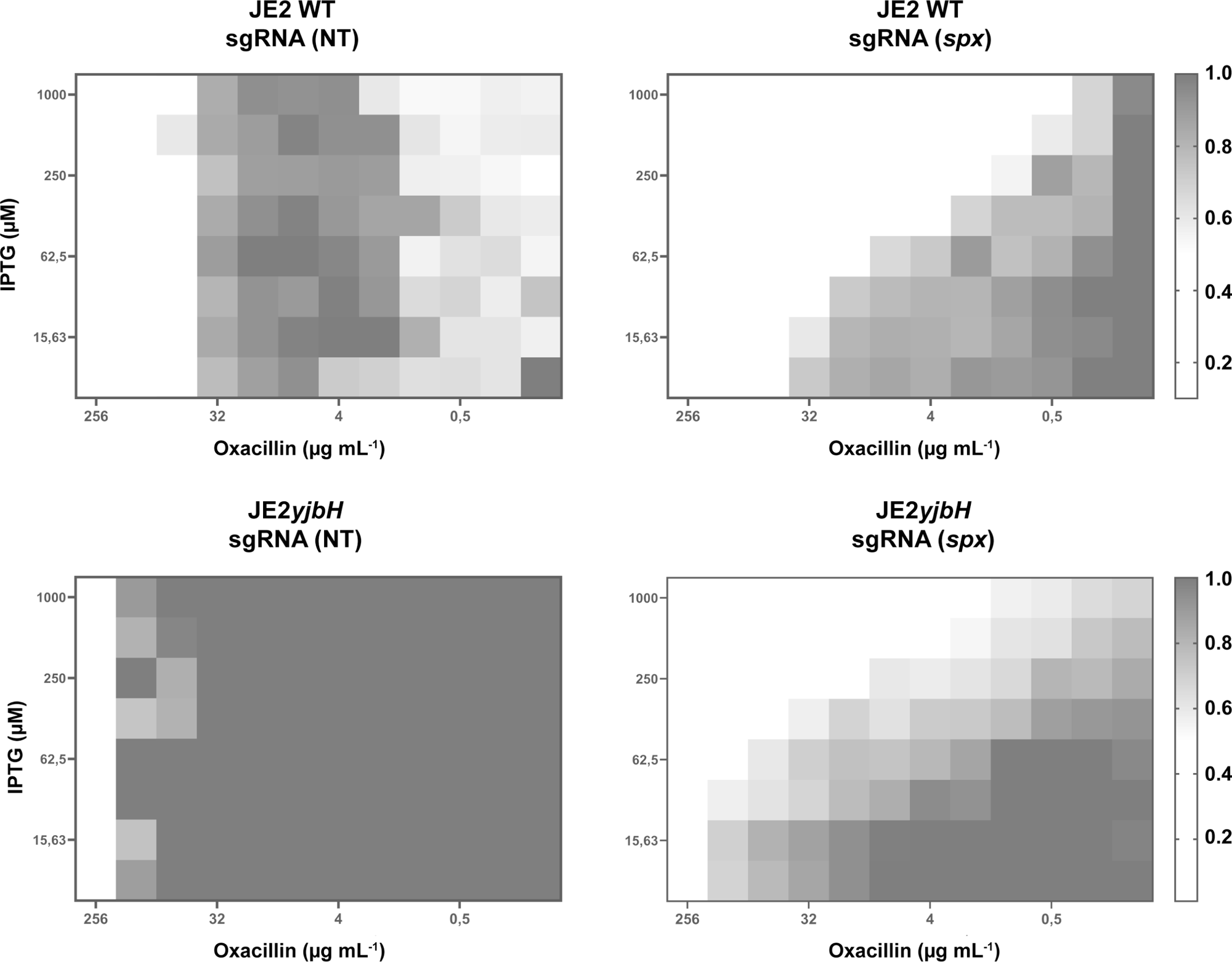
Oxacillin MICs correlates positively with increasing Spx levels. Checkerboard microdilution assays performed with increasing concentrations of oxacillin and increasing concentrations of IPTG to deplete Spx in the JE2 wild-type and in JE2 *yjbH* mutant. Plates were incubated at 37°C for 24 hours. The extent of inhibition is calculated as the OD_600_ relative to the untreated control (0 µM IPTG and 0 µg mL^-1^ oxacillin, lower right corner) and is shown as a heat plot.

### Spx is essential for growth at 30°C or in the presence of diamide

The results presented so far do not support the previous notion that Spx is essential for growth of *S. aureus* under standard laboratory conditions (39). None-the-less, multiple attempts to delete *spx* by allelic exchange in the USA300-JE2 strain were unsuccessful. Mutant construction by allelic replacement involves temperature shifts to non-optimal temperatures and we next asked if depletion of Spx confers a growth defect at temperature at 30°C or 42°C that may select against deletion of the gene. To answer this question, 10-fold serial dilutions of exponential JE2 wild-type and *yjbH* disrupted cells expressing either the NT-sgRNA or the *spx* specific sgRNA were spotted on plates with or without IPTG and incubated at 30°C, 37°C or 42°C for 24 hours. As shown in Supplemental Fig. 2A, cells depleted for Spx were unable to form visible colonies at 30°C while Spx depletion did not compromise colony formation at 37°C and 42°C.

The Spx transcriptional regulator is known to have a pivotal role in promoting growth during disulfide stress (25,41) and consistent with this role, depletion of Spx reduced the plating efficiency of JE2 cells approximately 1000-fold in the presence of 0.5 µM diamide (Supplemental Fig. 2B). At this concentration of diamide, the *spx* CRISPRi-strain conferred a growth defect to JE2 wild-type cells even when grown in the absence of IPTG (Supplemental Fig. 2B). Hence, the leakiness of the promoter controlling dCas9 transcription seems to result in sufficient Spx reduction to confer a growth defect, indicating that even a small reduction in Spx levels severely reduce fitness of wild-type cells in the presence of diamide (Supplemental Fig. 2B). Unexpectedly, cells with disrupted *yjbH* displayed decreased plating efficiency as compared to wild-type cells at 0.5 µM diamide (see cells grown in the absence of IPTG in Supplemental fig. 2B), showing that YjbH promotes growth of *S. aureus* during diamide-stress and that high Spx cannot compensate for YjbH. Moreover, knock-down of *spx* expression did not alter diamide-sensitivity in cells devoid of YjbH activity indicating that Spx depends on YjbH for promoting growth of *S. aureus* during disulfide stress (Supplemental Fig. 2B).

### High Spx broadly potentiates resistance to compounds targeting the cell envelope

Mutations that inactivate the YjbH-ClpXP protease complex responsible for Spx degradation have on several occasions been identified in clinical strains that developed resistance to these antibiotics during treatment (42–45). This prompted us to investigate if inactivation of *yjbH* or depletion of Spx would impact growth of *S. aureus* in the presence of these critical antibiotics. In spot-dilution assays, cells with inactivated *yjbH* expression showed improved plating efficiency (> 100 folds) in the presence of 1 µg ml^-1^ vancomycin, however, only under conditions allowing for expression of Spx (Fig. 5A). In the presence of 1 µg ml^-1^ vancomycin, the *spx* CRISPRi strain had reduced growth compared to wild-type cells even in the absence of IPTG, an observation that we ascribe to leakiness of the promoter controlling dCas9 expression (Fig. 5A). A similar picture was observed upon plating cells in the presence of 0.35 µg ml^-1^ daptomycin, where the *yjbH* cells showed improved plating efficiency and increased colony size under conditions allowing for Spx expression (Fig. 5A). As for vancomycin, the *spx* CRISPRi strain grown in the presence of 0.35 µg ml^-1^ daptomycin displayed reduced plating efficiency compared to the wild-type and the *yjbH* mutant even in the absence of IPTG (Fig. 5A). Together these data illustrate that the ability of *S. aureus* to grow in the presence of daptomycin or vancomycin correlates positively to Spx expression and that a small reduction in Spx levels (caused by the leaky expression of dCas9) confers hyper-sensitivity to both antibiotics.

**Fig. 5.**
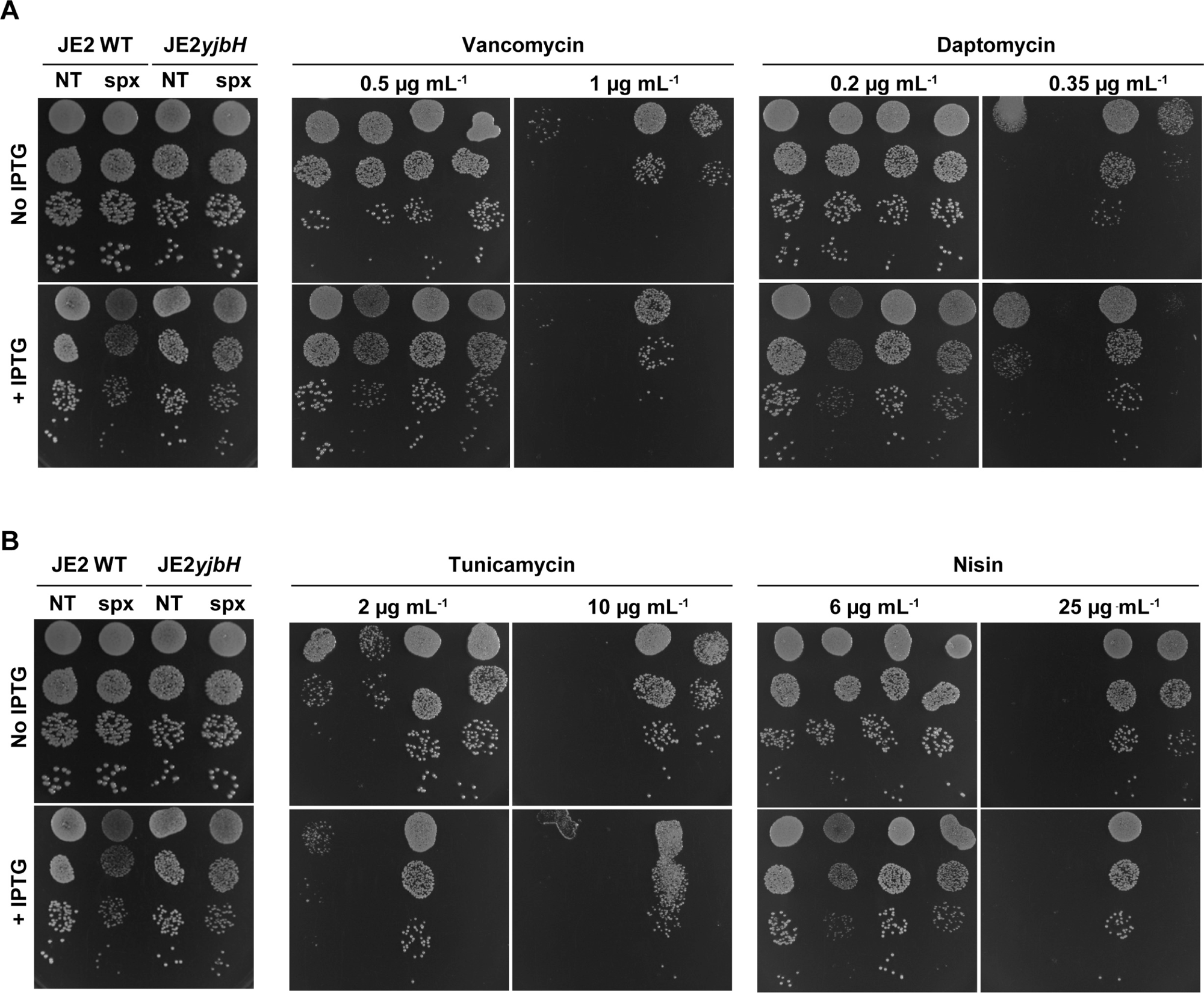
High Spx broadly potentiates resistance to compounds targeting the cell envelope. JE2 wild-type and JE2yjbH cells harboring the two plasmids allowing CRISPRi knock down of *spx* transcription were grown in the TSB at 37°C. At OD_600_ ∼ 0.4, ten-fold serial dilutions of each strain were spotted in 10 μL aliquots on TSA ± 1000 µM IPTG and with increasing concentrations of (A) vancomycin or daptomycin, or (B) tunicamycin or nisin as indicated. JE2 wild-type and JE2 *yjbH* mutant expressing a NT sgRNA were included as controls.

We proceeded by testing the susceptibility of the strains to compounds targeting other structures in the cell envelope: tunicamycin targeting the first step of the biosynthesis of the wall teichoic acid (WTA), an important constituent of the cell wall of Gram-positive bacteria, and nisin targeting the cytoplasmic membrane. Intriguingly, inactivation of *yjbH* allowed *S. aureus* cells to grow at greatly increased concentrations of tunicamycin and nisin, however, only if cells expressed Spx (Fig. 5B). Finally, we assessed if high Spx levels also increase resistance to two antibiotics with a primary target outside biosynthesis of the cell envelope, namely, ciprofloxacin that targets DNA replication, and tetracycline targeting protein synthesis. For these compounds inactivation of *yjbH* neither changed the MIC nor did it promote growth in spot-dilution assays (Supplementary Fig. 3; Supplementary Table 3). CRISPRi knock down of Spx expression, however, did compromise growth in the presence of ciprofloxacin (Supplementary Fig. 3). We conclude that high Spx specifically potentiates resistance to compounds targeting the cell envelope while reducing Spx levels may more broadly increase sensitivity to antibiotics.

### Increased Spx potentiates resistance during anaerobic growth and the potentiating effect is independent of the N-terminal redox sensing switch

There is a body of studies suggesting that reactive oxygen species (ROS) produced during respiration contribute to the killing mechanism of many types of antibiotics (reviewed in 46, 47). Hence, we speculated that Spx, an activator of an oxidative stress response, may potentiate antibiotic resistance by increasing the capacity of *S. aureus* cells to mitigate oxidative stress. To test this hypothesis, we repeated the spot-dilution assays, this time testing the ability of cells to grow in the presence of oxacillin, vancomycin, daptomycin, or nisin under anaerobic conditions (Supplementary Fig. 4). We first noted wild-type cells depleted for Spx formed smaller colonies than cells with normal levels of Spx in the absence of antibiotics suggesting that Spx also contributes to anaerobic growth (Supplementary Fig. 4). Importantly, anaerobic incubation did, however, not lead to major changes in the susceptibility of the strains to any of the compounds and, moreover, Spx-depleted cells were equally sensitive to oxacillin, vancomycin, and nisin when grown under anaerobic conditions (Supplementary Fig. 4). Only Spx-depleted cells spotted on plates with 0.35 µg ml^-1^ daptomycin seemed to have higher plating efficiencies on plates incubated anaerobically than on plates incubated with oxygen, suggesting that oxygen could play a role in *S. aureus* sensitivity to daptomycin (Supplementary Fig. 4).

Spx homologs have a conserved N-terminal redox-active C-X-X-C motif that upon oxidation forms an intramolecular disulfide bond that converts Spx to a transcriptional activator of oxidative stress genes (25,48). To further assess if a Spx-controlled oxidative stress response contributes to antibiotic resistance, we inactivated the redox-sensing C10-X-X-C13 motif of *S. aureus* Spx by substituting each of the two Cysteines with Alanine. While we were unable to delete the *spx* gene in JE2, JE2 strains harboring alleles encoding the Spx variants with substitutions in the redox sensing switch (Spx_C10A_, Spx_C13A_ or Spx_C10A + C13A_) were easily generated suggesting that inactivation of the C10-X-X-C13 motif, as opposed to deletion of *spx*, is not associated with a severe fitness-cost. Interestingly, inactivation of the redox switch also did not change the sensitivity of JE2 cells to oxacillin neither in spot-dilution assays (Fig. 6A) nor in microbroth dilution assays (Table 3).

**Fig. 6.**
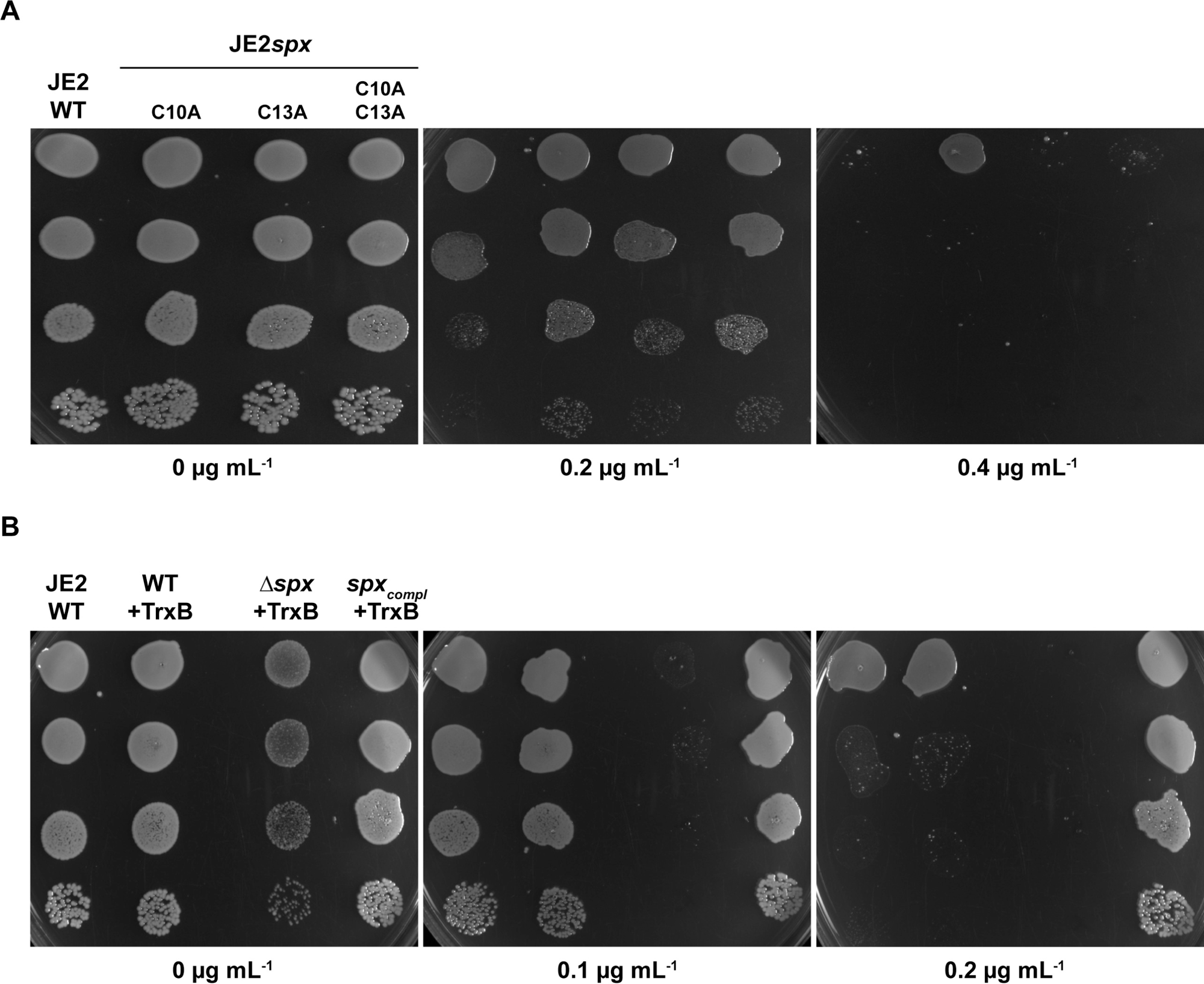
Spx potentiates resistance independently of the N-terminal redox sensing switch and overexpression of TrxB does not compensate for Spx in the presence of oxacillin. (A) The JE2 WT strain and JE harboring *spx* alleles that prevent formation of the disulfide bond involved in redox-sensing were grown in the TSB at 37°C. At OD_600_ ∼ 0.4, ten-fold serial dilutions of each strain were spotted in 10 μL aliquots on TSA with increasing concentrations of oxacillin and incubated at 37°C for 24 hours. (B) The indicated strains allowing for *sarA* promoter driven *trxB* overexpression from the chromosomally integrated pAQ69 plasmid were grown in TSB at 37oC. Spx was complemented by introducing a low-copy plasmid, pAQ75, expressing *spx* from its native promoter. At OD_600_ ∼0.4, ten-fold serial dilutions were spotted in 10 μL aliquots on TSA plates with increasing concentrations of oxacillin and were incubated at 37°C for 24 hours.

**Table 3.**
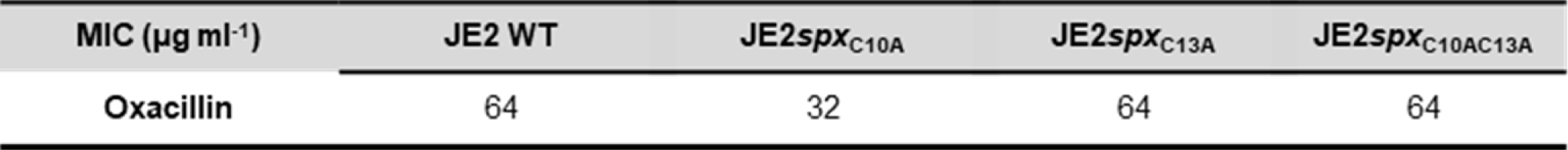
Oxacillin susceptibility of USA300-JE2 strain and *spx* derivatives with inactivated C-X-X-C motif.

| MIC ($\mu\text{g ml}^{-1}$) | JE2 WT | JE2*spx*<sub>C10A</sub> | JE2*spx*<sub>C13A</sub> | JE2*spx*<sub>C10AC13A</sub> |
| --- | --- | --- | --- | --- |
| Oxacillin | 64 | 32 | 64 | 64 |

Previous work showed that the essentiality of *spx* can be bypassed by overexpressing thioredoxin (TrxA) or thioredoxin reductase (TrxB) (39). Consistent with this notion, we were able to delete the *spx* gene in JE2 cells having the *trxB* overexpression plasmid pAQ69 integrated in the chromosome. When plated under standard conditions at 37°C, the JE2::pAQ69*Δspx* strain formed colonies of reduced size as compared to wild-type strain (Fig. 6B). Of particular importance to this study, the JE2::pAQ69*Δspx* strain was unable to form colonies at low concentrations of oxacillin demonstrating that overexpression of TrxB does not bypass the need for Spx in the presence of β-lactams antibiotics. On the other hand, introduction of the low copy number plasmid, pAQ75, carrying the wild-type *spx* gene under control of its native promoter control fully restored growth of JE2::pAQ69*Δspx* cells in the presence of oxacillin. Altogether, the results support that Spx controls resistance to compounds targeting the cell envelope by a mechanism that goes beyond the role of Spx in sensing and responding to oxidative stress.

## Discussion

The most used antibiotics target biosynthesis of the bacterial cell wall with the β-lactam class of antibiotics standing out as the most prescribed drug world-wide (49). Bacterial resistance to this group of antibiotics is, therefore, particularly challenging. In *S. aureus*, the selection and dissemination of the *mecA* gene carried by the staphylococcal cassette chromosome (SCC*mec*) element has been the main driver of resistance (5), however, here we identify the *yjbH* gene as a potentiator that when inactivated transforms the clinically important USA300 MRSA strains into being homogeneously highly resistant to β-lactam antibiotics. Inactivation of YjbH prevents Spx from being degraded by the ClpXP protease, and we further demonstrate that the accumulating Spx is causing the reduced susceptibility to β-lactams and other antibiotics targeting the cell envelope. Consistent with this finding, we previously showed that inactivation of the ClpXP protease confers hyper-resistance to β-lactams in MRSA belonging to the USA300 clone (12,21). However, the role of Spx in resistance was not addressed because the *spx* gene was claimed to be essential based on findings showing that i) the published *S. aureus* Δ*spx* contained a suppressor mutation in *rpoB* that seemingly compensated for the lethality of the *spx*-deletion, and ii) that an *spx* deletion could not be crossed into wild-type *S. aureus* strains using generalized bacteriophage-mediated transduction (39,41). Thus, our study illustrates the value of CRISPRi for studying the essentiality of genes under different growth conditions. In a few cases, accumulation of Spx has been experimentally verified in clinical MRSA strains that developed resistance during treatment (50). The clinical relevance of Spx stabilization for *S. aureus* survival during antibiotic treatment is further supported by the selection of mutations that inactivate the YjbH-ClpXP protease in *S. aureus* isolates from patients undergoing treatment with β-lactams, daptomycin or vancomycin (42–45, 51). Moreover, a comprehensive genetic study identified the *yjbH* gene as a hot spot for adaptive mutations during infections supporting that *S. aureus* may benefit from high Spx levels in clinical settings, but the role of antibiotics in selection was not discussed in this study (28). Low-level oxacillin resistance can arise in *mecA-* or *mecC*-negative *S. aureus* isolates, and such isolates are often referred to as <u>b</u>orderline <u>o</u>xacillin-resistant *<u>S</u>. <u>a</u>ureus* (BORSA) (52). The BORSA phenotype has traditionally been attributed to mechanisms involving hyper production of β-lactamase, mutations in native PBPs, or elevated PBP4 levels (32,53–54). Interestingly, mutations in *yjbH* were previously associated with non-*mec*-mediated oxacillin resistance and overproduction of PBP4, (29–32, 34). In our study, we confirmed that inactivation of *yjbH* was associated with a slight upregulation of PBP4, however, with the CRISPRi tool in hand we could show that elevated levels of PBP4 did not contribute to the resistant phenotype and that high Spx even bypasses the role of PBP4 in resistance. Of note, a comprehensive genomic study recently identified the *clpX* gene as a hot-spot for adaptive mutations that drive non-mec-mediated oxacillin resistance (51). Therefore, Spx stabilization may be a common pathway that potentiates *S. aureus* β-lactam resistance in strains with or without *mecA,* necessitating further investigation of this clinically important phenotype.

Oxidative stress induces formation of a disulfide bond in Spx that activates it for promoting transcription of a disulfide stress response (25,48). Our data suggested that Spx potentiates antibiotic resistance independently of its redox-sensing cysteines. Consistent with our data, Göhring et al (29) found that while the four cysteine residues of *S. aureus* YjbH were essential for its ability to affect susceptibility to disulfide stress, they were dispensable for the effect on β-lactam susceptibility in methicillin sensitive *S. aureus*. Hence, the activation of Spx in disulfide stress and β-lactam resistance appear to rely on different types of mechanisms. Of high relevance, Spx was recently shown to be stabilized by cell wall stress in the soil-bacterium *Bacillus subtilis* (55,56). Stabilization is mediated by the anti-adaptor protein YirB, which by binding to YjbH protects Spx from being degraded by ClpXP (56). Interestingly, under cell wall stress, *B. subtilis* Spx remains in the reduced state supporting that the redox-active disulfide switch is dispensable for the activation of Spx during cell wall stress (55,56). When we performed a BLAST search, we did not identify homologs of YirB in the S*. aureus* proteome. The Renzoni group, however, observed that the levels of Spx go up in *S. aureus* cells exposed to oxacillin and vancomycin, and that Spx stabilization was induced by aggregation of YjbH (57). In contrast, ribosome-targeting antibiotics, kanamycin, tetracycline, and erythromycin treatments did not increase total levels of Spx in *S. aureus* (57). Altogether, these findings led us to hypothesize that during conditions of cell wall stress, the *S. aureus* Spx regulator is stabilized to activate transcription of a more general stress response that broadly protects MRSA from antibiotics targeting the cell envelope. The inactivation of YjbH was previously demonstrated to induce major changes in the transcriptome of non-stressed *S. aureus* cells, with the genes encoding proteins involved in urea uptake and degradation being strongly upregulated while surface-associated virulence factors such as the immunoglobulin G binding protein A and Sbi and genes encoding extracellular proteases, lipases, and thermos-nuclease being strongly down-regulated (58). Likewise, ClpXP inactivation resulted in a strong upregulation of the urease operon and strong downregulation of the listed virulence factors (59). This striking overlap in the transcriptional changes induced by inactivation of ClpXP or YjbH indicates that the observed changes are attributable to stabilization of Spx. Future studies will be directed towards understanding how Spx controlled genes potentiate *S. aureus* resistance to compounds targeting the cell wall. Of high relevance, disruption of the *yjbH* gene was previously demonstrated to induce large changes in the composition of cell wall-glycopolymers. Spx does not control gene-expression through direct binding to DNA (25). Instead, Spx changes transcription through direct binding to the RNA polymerase subunits (25). Strikingly, mutations in the RNA polymerase subunits RpoB and RpoC are the most frequently encountered potentiators of antibiotic resistance in *S. aureus* and we hypothesize that potentiating mutations in the RNA polymerase and high Spx upregulate the same set of genes.

In conclusion, our study leads support to a novel paradigm claiming that development of high-level resistance in MRSA is a two-step process where the introduction of *mecA* in itself only results in a modest increase in resistance that is displayed in a heterogeneous manner, and that the development of high-level resistance is associated with genetic adjustments in potentiator genes. Bacterial cell wall synthesis takes place in the outer wall and at the division septa and requires the coordinated activities of many proteins working together in machines. We speculate that the Spx induced stress response mitigates the cell wall stress elicited by integrating PBP2A, which has evolved outside of *S. aureus*, into the native cell wall synthesizing apparatus. Potentiators therefore represent an Achilles heel in bacterial resistance development that could pave the way for novel, innovative therapeutic approaches that if targeted could suppress bacterial resistance.

## Materials and methods

### Strains and culture conditions

The strains used in this study are listed in Table S1. Unless stated otherwise, *S. aureus* strains were cultivated in tryptic soy broth media (TSB; Oxoid) under agitation at with linear shaking at 180 rpm at 37°C. For solid medium, 1.5% agar was added to make TSA plates. The growth was assessed by measuring optical density at a wavelength of 600 nm (OD_600_). In all experiments, we used bacterial strains freshly streaked from the frozen stocks on TSA plates and incubated overnight at 37°C. In most experiments, 20 mL of TSB culture was inoculated in 200-mL flasks to allow efficient aeration of the medium with a starting OD of < 0.05. Antibiotics were added as indicated. The strains containing the pLOW-dCas9_aad9 and pVL2336-sgRNA plasmids for the CRISPRi system were cultured or plated with 250 µg mL^-1^ spectinomycin and 10 µg mL^-1^ chloramphenicol. For induction of dCas9 expression, 1000 µM isopropyl *β*-D-1-thiogalactopyranoside (IPTG) was added unless stated otherwise. *E. coli* strains were grown in LB, ampicillin was added at 100 µg mL^-1^.

#### Spot dilution assay

Freshly streaked *S. aureus* strains were inoculated in TSB, supplied with antibiotics if required, and the cultures were grown at 37°C with aeration and shaking at 180 rpm until early exponential phase growth (OD_600_ = ∼ 0.5). Tenfold serial dilutions were prepared in 0.9% NaCl solution and 8-10 µL aliquots of each dilution was spotted onto TSA plates containing appropriate antibiotics. Plates were incubated overnight at the appropriate temperature. Spot dilution assays were performed twice in triplicates to ensure reproducibility

#### Growth curves

Growth curves were obtained at 37°C with continuous orbital shaking by following optical density (OD_600_) in 96-well microtiter dishes using the Synergy H1 Multimode Reader (BioTek^®^). OD_600_ was measured every 20 min, and the values were log_10_-transformed in the graphs. Growth assays were performed twice in triplicates to ensure reproducibility.

### Construction of CRISPRi knock down mutants

The CRISPRi plasmids used in this study are described in Liu et al., (60). Transformation in *E. coli* IM08B and *S. aureus* JE2 was done as described herein (60). Uptake of the pLOW-dCas9_aad9 plasmid was confirmed by PCR with primers pLOW_aad9_F and pLOW_aad9_R. The No-target sgRNA and sgRNA specific for *pbp4* and *spx* were cloned into vector pVL2336 using Golden Gate cloning as described in Liu et al. (60). The purified pVL2336-sgRNA plasmids were transformed into *S. aureus* JE2 and successful plasmid transformation was confirmed by PCR using the primers pVL2336_5517bp_F and pVL2336_339bp_R. All Primers used in this study are listed in Table S2.

### Construction of *spx* mutants

The *S. aureus* JE2 Δ*spx* in-frame marker less gene deletion mutant, along with the *spx*_C10A_, *spx*_C13A,_ and *spx*_C10,13A_ point mutants, were generated using pJB38 plasmid derivatives: pAQ5, pAQ25, pAQ43, and pAQ42, respectively (61). All pJB38 plasmid derivatives were assembled using NEBuilder high-fidelity DNA assembly cloning kit according to the manufacturer’s instructions. Primers used for amplifying DNA fragments containing the engineered mutations for construction of pAQ5, pAQ25, pAQ43 and pAQ42 are listed in Table S2. The allelic exchange process with pJB38 derivatives was carried out as described recently (61). The Δ*spx* mutation was constructed in *S. aureus* JE2 with pAQ69 integrated into the SAPI chromosomal site. The plasmid pAQ69 was constructed by PCR amplifying P*_sarA_*-*trxB* DNA sequence from pAQ24 and cloning into the multiple cloning site of pJC1111, a SAPI integration vector. The pAQ24 plasmid was constructed by replacing the dsRED allele in pVT1 (62) with *trxB* amplified from JE2 genomic DNA. The integration of pAQ69 into the SAPI site of *S. aureus* JE2 was performed as previously described (63). The plasmid pAQ75 used to complement the Δ*spx*::pAQ69 mutant was constructed by amplifying *spx* along with its native promoter and cloning into pKK22, a stable low copy vector for *S. aureus* (64). The primers used to amplify DNA segments for pAQ69, pAQ75 and pAQ24 are listed in Table S2 and the plasmids were assembled using the NEBuilder high-fidelity DNA assembly cloning kit.

#### Minimum Inhibitory Concentration

Minimum Inhibitory concentration (MIC) was determined by following the Clinical and Laboratory Standards Institute 2017 guidelines in the 96-well format. Overnight cultures of *S. aureus* were diluted in physiological saline (0.9% NaCl) to reach turbidity of 0.5 McFarland (Sensititre^®^ nephelometer and the Sensititre^®^ McFarland Standard). The bacterial suspensions were adjusted 5 x 10^5^ CFU mL^-1^ in cation-adjusted Mueller-Hinton broth in wells containing standard two-fold dilutions of the test antibiotics in a final volume of 100 µL. The plates were incubated for 16-20 h with low shaking at 37°C. When using *S. aureus* strains containing the CRISPRi system, 250 µg mL^-1^ spectinomycin and 10 µg mL^-1^ chloramphenicol was added while 1000 µM IPTG was added when indicated. MIC was defined as the lowest concentration of the antibiotic at which visible growth was completely inhibited. MIC assays were performed in biological duplicates to ensure reproducibility.

#### Population analysis profiles

Antibiotic resistance profiles were determined as previously described (50). Overnight cultures of tested strains were normalized to an OD_600_ of 1.0 and serially diluted to 10^-7^ in 0.9% NaCl. 10 µL of appropriate dilutions were spotted on TSA plates supplemented with increasing concentrations of oxacillin. The number of CFU mL^-1^ was determined after 24 h of growth at 37°C. The data shown is representative for two biological duplicates. Population analysis profiles were performed in biological triplicates to ensure reproducibility.

#### Checkerboard analysis

Oxacillin and IPTG were 2-fold serially diluted at ten and eight different concentrations, respectively, to create a 10 x 8 matrix (see supplemental Figure 1). Stock solutions of oxacillin (256 – 0.25 µg mL^-1^) and IPTG (1000 – 15.63 µM mL^-1^) were prepared in 0.9% NaCl and aliquots of 20 µL were added to the 96-well plate. Overnight cultures of *S. aureus* were diluted in 0.9% NaCl to reach turbidity of 0.5 McFarland (Sensititre^®^ McFarland Standard) and the bacterial suspensions were adjusted 5 x 10^5^ CFU mL^-1^ in TSB with 250 µg mL^-1^ spectinomycin and 10 µg mL^-1^ chloramphenicol to retain the CRISPRi system. 180 µL aliquots were dispensed into all wells. Plates were incubated at 37°C for 20-24 h. The experiment was performed in three biological triplicates.

#### Western blot analysis

To determine the levels of PBP2a, Spx, and PBP4 in *S. aureus* cells, the bacterial strains were grown in TSB at 37°C with aeration until OD_600_ reached 1.0. Once this point was reached, 1 mL of cells from each tested strains was harvested, and an extract of total cellular proteins was prepared by harvesting the cells by centrifugation and resuspending cell pellets in 50 mM Tris-HCl (pH 8.0) (200 µL per OD unit) and incubated with 5 µg mL^-1^ lysostaphin (Sigma) for 30 min at 37°C. To determine the amount of PBP2a in the cells of *S. aureus* grown in the absence or presence of oxacillin, exponentially growing cultures (OD_600_ = 0.8) were divided into two cultures that continued growth in the absence or presence of 0.2 µg mL^-1^ oxacillin for an additional 30 min. Total cellular proteins were purified from 1 mL of culture as described above. 20 µL of each sample was loaded on NuPAGE 4 to 12% Bris-Tris gels (Invitrogen), and electrophoresis was performed according to the manufacturer’s instructions. After separation, proteins were blotted onto a polyvinylidene difluoride (PVDF) membrane (Invitrogen) using an xCell II blot module system (Invitrogen). To detect PBP2a, the PVDF membrane was first blocked with human IgG to block protein A. PBP2a was detected using mouse anti-staphylococcal PBP2a (Merck) at a 1:2.500 dilution. PBP4 was probed using rabbit anti-staphylococcal PBP4 (a generous gift from Professor Mariana Pinho, Portugal) at a 1:2.500 dilution. The Spx was probed using rabbit anti-*Bacillus subtilis* Spx at a 1:2.500 dilution (kindly provided by Professor Peter Zuber, Oregon, US). Detection of the specific protein signal was achieved by using WesternBreeze Chemiluminescent (anti-rabbit or anti-mouse) kit (Invitrogen) or by using Immobilon Forte Western HRP substrate (Merck)

## Figure Legends

**Supplemental Fig. 1. Overview of the CRISPRi system and MIC setup.** (A) the pLOW-dCas9_aad9 carries the *dcas9* gene under the control of an IPTG-inducible promoter while the sgRNA is constitutively expressed from the pVL2336-sgRNA. In the presence of IPTG, the combined expression of dCas9 and the gene specific sgRNA causes a transcription block of the gene of interest. (B) Illustration of sgRNA annealing sites in the *pbp4*- and *spx* genes. (C) Setup of MIC assays conducted with strains carrying the CRISPRi system. The MIC assay was conducted in TSB ± IPTG as indicated and with the strains expressing either a gene specific sg-RNA or a non-targeting RNA.

**Supplemental Fig. 2. Depletion of Spx confers a growth defect at 30°C and during disulfide stress.** The JE2 WT strain and the JE2 yjbH mutant expressing the CRISPRi system with no-target (NT) or *spx* sgRNA were grown in TSB at 37°C with shaking. At OD_600_ ∼ 0.4, ten-fold serial dilutions of each strain were spotted in 10 μL aliquots on TSA ± 1000 µM IPTG that were incubated either (A) at different temperatures for 24 hours, or (B) at 37°C for 24 hours with increasing concentrations of diamide.

**Supplemental Fig. 3 High Spx levels do not increase resistance towards tetracycline or ciprofloxacin.** (A) JE2 WT and JE2yjbH expressing the CRISPRi system with no-target (NT) or *spx* targeting sgRNA were grown in TSB supplied with appropriate antibiotics at 37°C. Once exponential growth was reached, 10 µL aliquots of 10-fold serial dilutions were spotted on TSA in the absence (0 µM) or presence (1000 µM) of IPTG supplied with increasing concentrations of tetracycline and ciprofloxacin.

**Supplemental Fig. 4. Spx depletion confers hyper-sensitivity to antibiotics targeting the cell envelope also under anaerobic conditions.** JE2 WT and JE2 yjbH expressing the CRISPRi system with no-target (NT) or *spx* targeting sgRNA were grown in TSB supplied with appropriate antibiotic at 37°C. Once exponential growth was reached, 10 µL aliquots of 10-fold serial dilutions were spotted on TSA in the absence (0 µM) or presence (1000 µM) of IPTG supplied with increasing concentrations of indicated antibiotics and set to incubate at 37°C for 24 h under aerobic or anaerobic (5% CO_2_) conditions.

## Acknowledgments

We greatly acknowledge A very special thanks to the staff at the Core Facility for Integrated Microscopy (University of Copenhagen) for their enthusiastic assistance in doing microscopy.

---

**Table S1.**
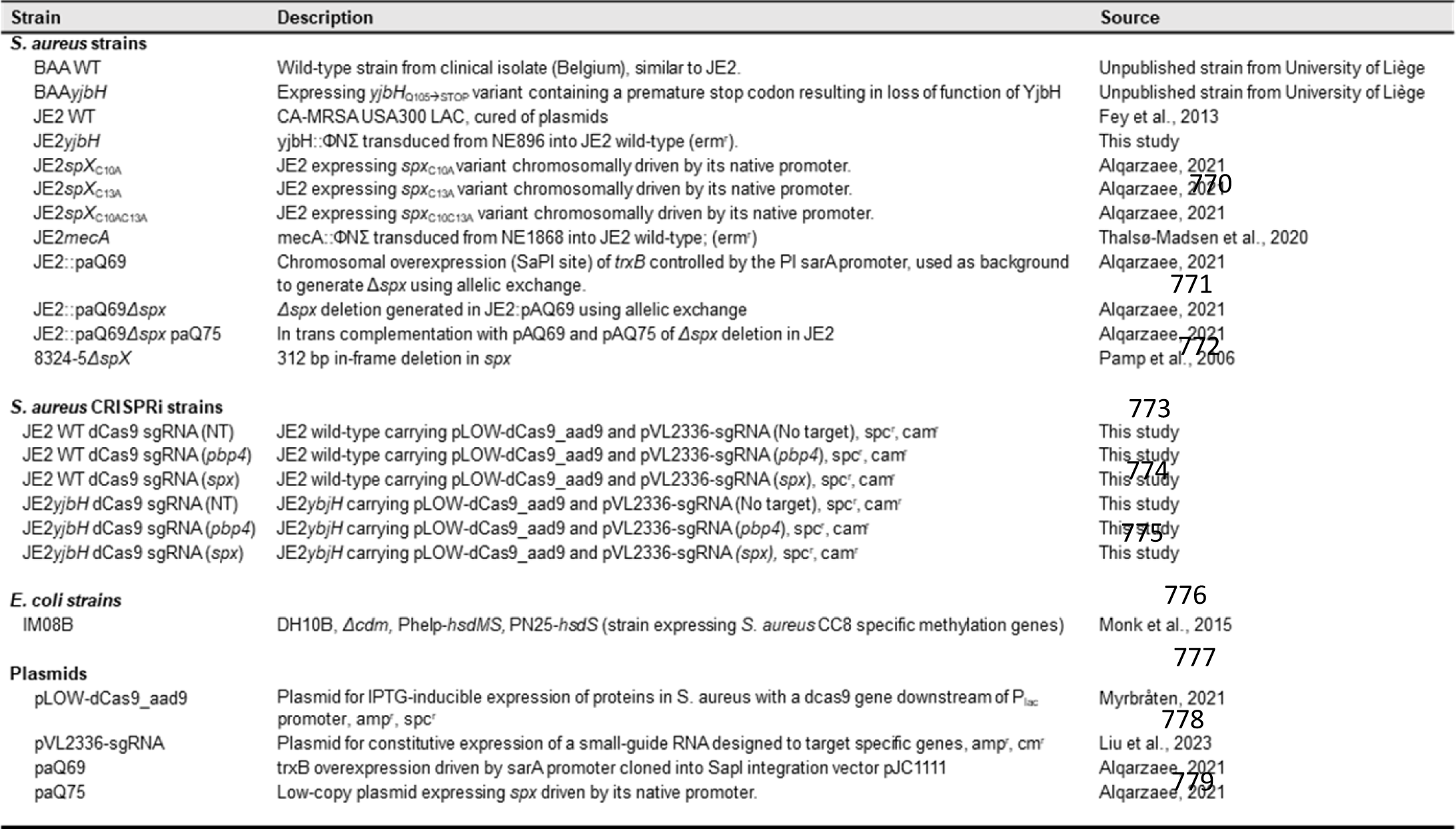
Bacterial strains and plasmids used in this study.

**Table S2.**
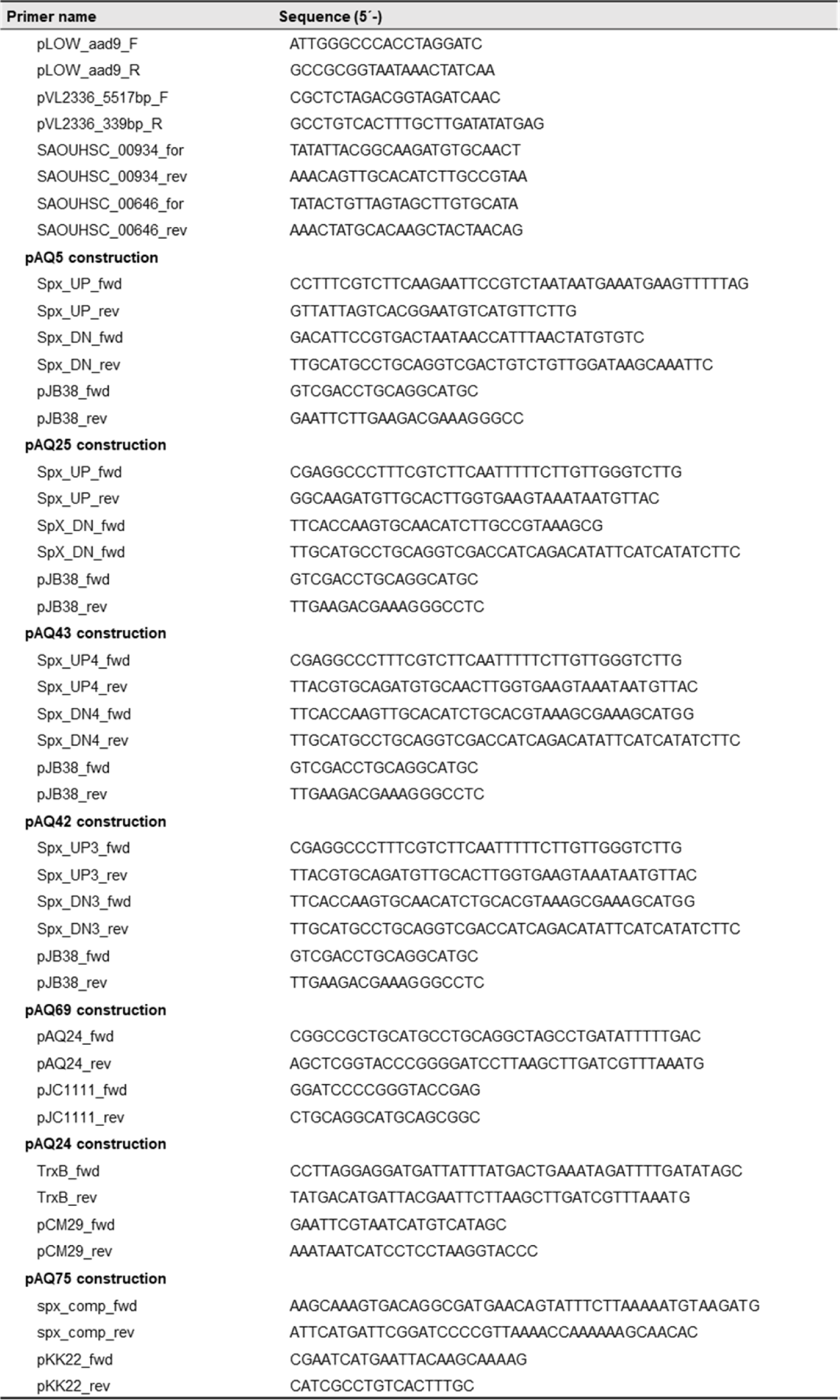
Primers used in this study.

**Table S3.**
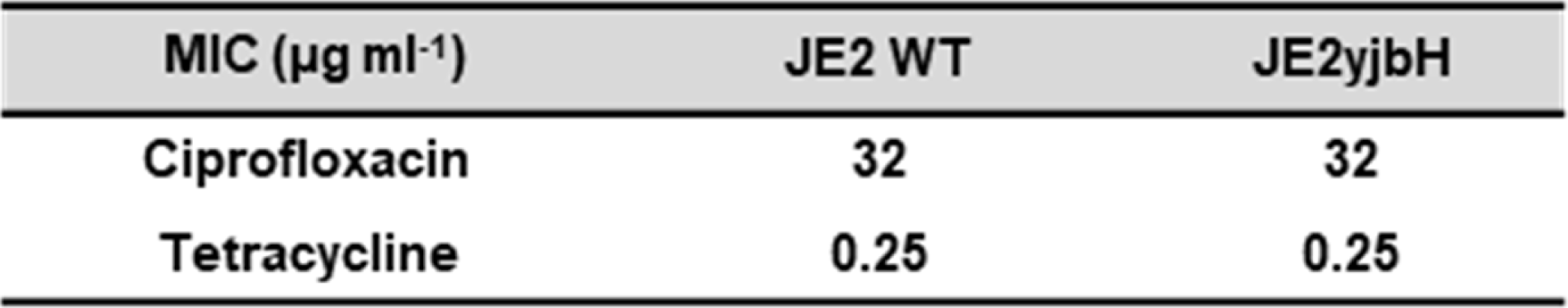
Antibiotic susceptibility of USA300-JE2 and the corresponding *yjbH* derivative to tetracycline and ciprofloxacin.

| MIC ($\mu\text{g ml}^{-1}$) | JE2 WT | JE2yjbH |
| --- | --- | --- |
| Ciprofloxacin | 32 | 32 |
| Tetracycline | 0.25 | 0.25 |

